# Genomic and Environmental Contributions to Chronic Diseases in Urban Populations

**DOI:** 10.1101/099770

**Authors:** Marie-Julie Favé, Fabien C. Lamaze, Alan Hodgkinson, Héloïse Gauvin, Vanessa Bruat, Jean-Christophe Grenier, Elias Gbeha, Kimberly Skead, Audrey Smargiassi, Markey Johnson, Youssef Idaghdour, Philip Awadalla

## Abstract

Uncovering the interaction between genomes and the environment is a principal challenge of modern genomics and preventive medicine. While theoretical models are well defined, little is known of the GxE interactions in humans. We used a system biology approach to comprehensively assess the interactions between 1.6 million environmental exposure data, health, and expression phenotypes, together with whole genome genetic variation, for ∼1000 individuals from a founder-population in Quebec. We reveal a substantial impact of the urbanization gradient on the transcriptome and clinical endophenotypes, overpowering that of genetic ancestry. In detail, air pollution impacts gene expression and pathways affecting cardio-metabolic and respiratory traits when controlling for genetic ancestry. Finally, we capture 34 clinically associated expression quantitative trait loci that interact with the environment (air pollution). Our findings demonstrate how the local environment directly affects chronic disease development, and that genetic variation, including rare variants, can modulate individual’s response to environmental challenges.

**Highlights:**

- Fine scale environmental effects overpower those of ancestry on gene expression
- Air pollution (geographic and temporal) is associated with transcriptional response
- Gene-by-environment interactions with air pollution include asthma associated loci
- Inflammatory pathways and cardio-respiratory clinical traits are among those affected

## Introduction

Environmental exposures, together with genetic variation, influence disease susceptibility and elucidating their respective contributions remains one of the principal challenges of understanding complex diseases (*1*–*7*). Individuals with different genotypes may respond differently to environmental variation and generate different phenotypes (*8*–*11*). Such gene-by-environment interactions are thought to be pervasive and may be responsible for a large fraction of the unexplained variance in heritability and disease risk (*9, 12, 13*). Yet, disease risk, owing to either environmental exposures and/or their interactions with the genotype, remains poorly understood (*1, 2, 14*).

Canada’s precision medicine initiative, the Canadian Partnership for Tomorrow Project (CPTP (http://www.partnershipfortomorrow.ca)) is a cohort of over 300,000 Canadians, and capturing over 700 variables of longitudinal health information and environmental exposures to determine genetic and environmental factors contributing to chronic diseases. The program includes the Quebec regional cohort, CARTaGENE, having enrolled over 40,000, predominantly French-Canadian individuals between 40 to 70 years of age (*15*–*17*). Using individuals from this founding population, we selected 1007 individuals to determine how the genomes, the environment, and their interactions contribute to phenotypic variation. After attributing a regional and/or continental ancestry to each individual using genome-wide polymorphism data, we first captured the effect of different environmental exposures on gene expression and health related traits, while controlling for genetic relatedness and migration. Furthermore, to capture gene-by-environment interactions through eQTL analyses, we combined whole-transcriptome RNA-Sequencing profiles with whole-genome genotyping and rich fine-scale environmental exposure data.

Individuals living across a North-South urbanization cline in Quebec were selected for analyses, including: Montreal, the largest urban center in the Quebec province (MTL, 4500 individuals/km^2^); Quebec City, a smaller urban center (QUE, 1140 ind/km^2^); and Saguenay-Lac-Saint-Jean, a less urbanized region (SAG, 800 ind/km^2^). Differences in the regional environment within and across these cities, including ambient pollutant concentrations, have been associated with various health outcomes (*18, 19*). The majority of the Quebec population is of French-Canadian (FC) descent; a group of individuals descending from the French settlers that colonized the Saint-Lawrence Valley from 1608 to the British conquest of 1759 (*20, 21*). Despite considerable expansion, the population remained linguistically and religiously isolated while remote regions were colonized by small numbers of settlers, such as SAG (*22*, *23*) and contributed to the establishment of subpopulations. These sequential population bottlenecks impacted the genome of FCs by increasing the relative deleterious mutations load (*24*), while reducing overall genetic diversity in the population relative to the European population (*25*).

Using high-density whole-genome genotyping assays (Illumina Omni 2.5), and a combination of a haplotype-based methods powerful enough to detect fine-scale genetic structure (*26*) and data for country of birth for grand-parents, French-Canadians form a distinct genetic cluster relative to those of European descent (Fig. 1A, Fig SS1A to C), as has been previously observed (*24*). Within this FC group, we captured fine-scale regional genetic variation that traces a North-South cline across Quebec (Fig. 1B,C and Fig SS1D), consistent with Quebec settlement history and local ancestry. In addition to FCs, we identified 136 and 172 individuals of European and other ancestries respectively, reflecting recent immigration and admixture in Quebec (Fig SS1A to C).

**Figure 1:**
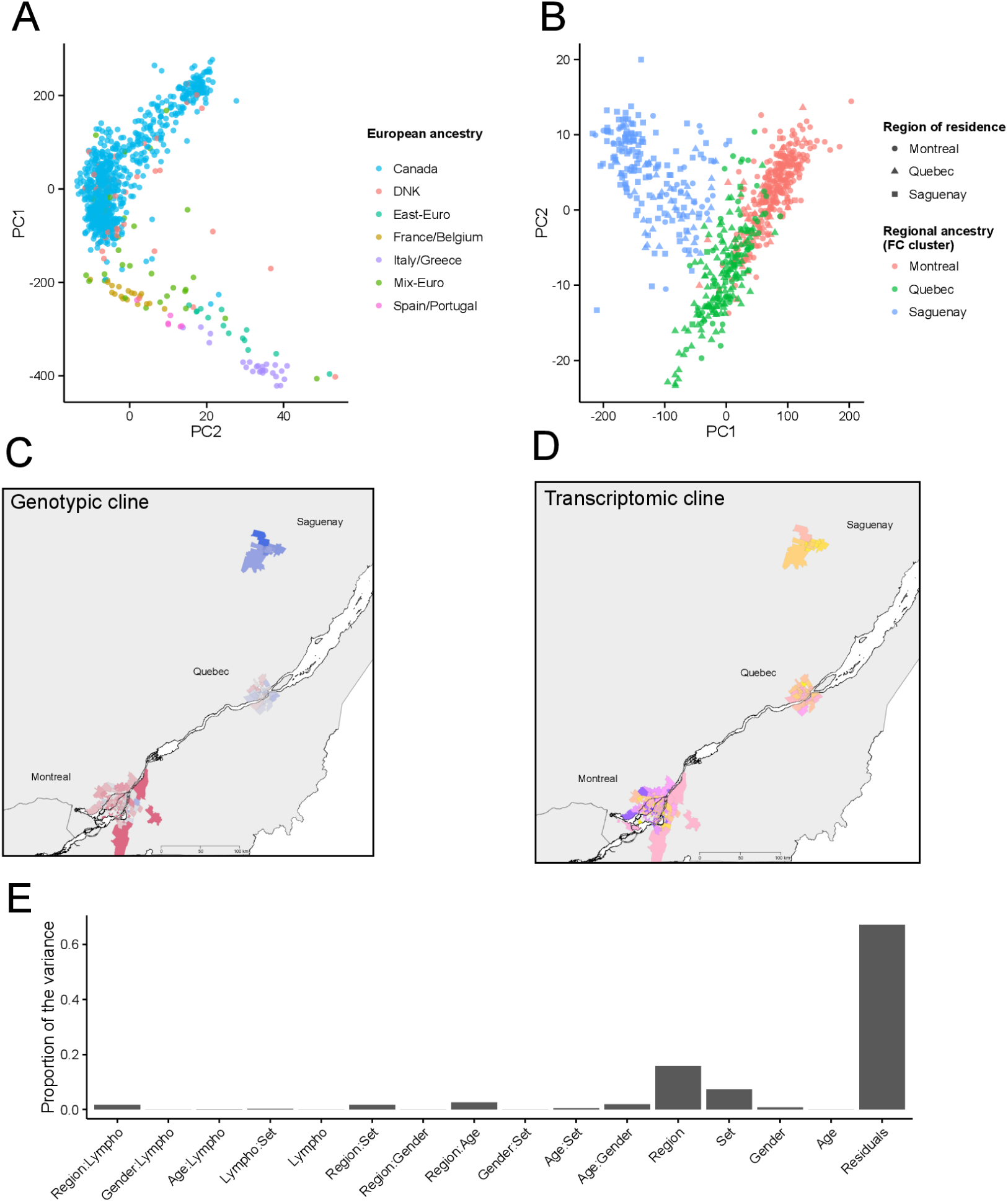
Genetic and transcriptomic variation within the CARTaGENE cohort sample. (A) Principal component analysis (PCA) of individuals of European descent, including FCs (n=887). Individuals are labeled according to self-declared ancestry based on the origin of four grand-parents.(B) PCA on the haplotype chunk (26) count matrix of French-Canadians (n=689) reveals 3 groups corresponding the region of residence, with SAG individuals showing less overlap with either of MTL or QUE individuals, in line with their historical isolation (*22, 23*). (C) Genotypic cline for individuals by location of residence (three-digit postal code) sampled across the province. Colour indicate the average value of the first principal component from a PCA on genotypes in each FSA (n=157). (D) Transcriptomic cline for individuals by location of residence (three-digit postal code) sampled across the province. Colours represent the average value of the first principal component from a PCA on the transcriptome in each FSA (n=189). (E) Proportion of transcriptomic variance (PVCA) in FCs explained by low-level phenotypes and their interactions.

High-coverage RNA-sequencing (approximately 60 million reads per individual) of all individuals reveals a similar geographic cline in transcriptional profiling (Fig. 1C). Geography (region of residence) and neutrophil counts each explain a significant proportion of the genome-wide transcriptional variation (Table S1). Using a test dataset of 708 individuals, we quantified the proportion of the variance in expression attributable to cell counts, age, sex, region, and arterial stiffness (See Supplemental material) by using principal variance component analysis (PVCA), and found that the region of residence explains ∼16% of the variance in gene expression, while the effects of age, sex, and cell counts were much lower (Fig. 1E). These analyses were repeated on an additional 289 participants and both of these effects were found to be replicated on expression profiles (Table S1). Similarly, when combining transcriptional profiles for all individuals, we found that the region of residence explains ∼15% of the variance in gene expression both in FCs and in Europeans (Fig. S2). Gene expression variation is not associated with the sampling clinic within a region (Fig. S3), and we applied stringent corrections that effectively remove batch effects (Fig. S4 and Table S1) therefore controlling for possible unwanted technical or biological variation in the differential expression analyses (see Supplemental materials for methodological details).

Given our ability to define FC individuals’ regional ancestry using whole-genome genotyping, we were able to ask whether population ancestry or regional environmental explains more of the transcriptional variation in the population. We first distinguished between “FC-locals” and “FC-internal migrants”; locals are FCs who have a regional genetic ancestry that is identical to the region where they currently reside, and internal migrants are FCs who have a regional genetic ancestry that differs from the region where they currently reside (Fig. 1B, Fig. S1D, Table S2). First, among FC-locals living along the North-South cline, we identified an increasing number of differentially expressed genes (DEG) and found that 505 significant DEGs (*p*-value < 0.05/15632, log-fold change (LFC) > 0.5) between Mtl-locals and Que-locals, 2167 between Que-locals and Sag-locals, and 6649 between Mtl-locals and Sag-locals (Fig. 2A). Additionally, we identified large numbers of DEGs between individuals with the same regional ancestry but who reside in different regions (FC-locals vs FC-internal migrants with the same genetic ancestry, but residing in different regions), and we find this pattern in nearly all pairwise com-parisons of this nature (Fig. 2B). On the other hand, when we performed comparisons between FC-locals and FC-internal migrants individuals who reside in the same region, but whose ancestry is from different parts of Quebec, we found very few DEGs in nearly all such comparisons (Fig. 2C).

**Figure 2:**
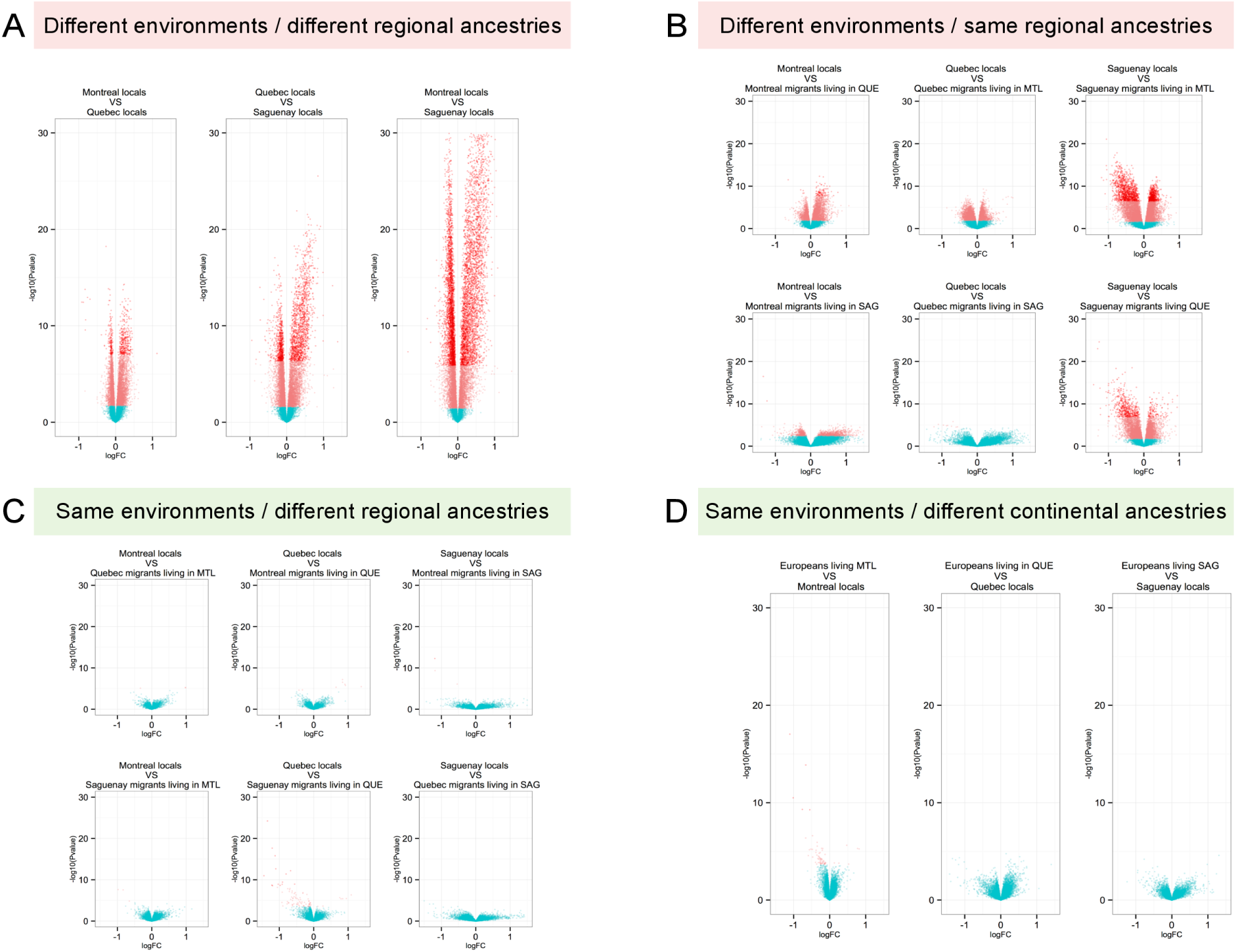
Environmental impacts on gene expression profiles override that of genotype. Contrasting the effects of ancestry and regional environment on DGE; (A) between FC-locals (different regional ancestry, different regional environments). (B) between FC-locals and FC internal migrants (same regional ancestry, different regional environments). (C) between FC-locals and FC internal migrants (different regional ancestries, same regional environment). (D) between FC-locals and Europeans (different continental ancestries, same regional environment). Pink dots are genes with FDR below 5% and Red dots are genes with p-value < bonferroni corrected p-value (3.20 x 10^−6^)

We replicated these findings through comparison of Europeans and FC-locals residing in the same region and found very few DEGs between them (Fig. 2D, Fig. S5). The lack of DEG is not due to differences in statistical power as we are able to identify up to 75% of our DEGs using only 30% of our FC individuals (Fig. S6). DEGs between regions are enriched for genes implicated in oxygen and gas exchange, G-protein coupled receptors, and inflammatory response (Fig. S7, Table S3). Although we initially captured both genotypic and transcriptional variation correlated with geographic clines among the French-Canadian population, these results indicate that shared regional environmental exposures influence peripheral blood expression profiles to a greater extent than regional or local (and continental) ancestry, and point to potential critical exposures contributing to pathways, phenotypic variation, and possibly disease development.

To test whether environmental exposures contribute to the geographic variation associated with transcriptional profiles and clinically relevant phenotypes across Quebec, we collated finescale environmental data (Fig. S8 and Fig. S9), including socio-economic indices, annual ambient air pollutant levels, vegetation indices (greenness), and “built environment” at the individual level for a total of 12 environmental exposures, at the scale of mail sortation area (these areas include several houses or a neighborhood, and their sizes are inversely proportional to population density - see Supplemental material). Indeed, these environmental exposures capture broad environmental correlates and variance across the Quebec province (Fig. S9), while allowing us to analytically treat individual specific exposures and ignore broad geographic sampling categories.

We found that the expression profiles of DEGs between regions, and the genes that regulate them (RDEGs), are largely associated with gradients of annual ambient air composition across Quebec (Fig. 3, Fig. S9). The North-South urbanization cline is indeed reflected by higher concentrations of particulate matter 2.5 (PM2.5) and nitrogen dioxide (NO2) in downtown MTL, however, higher concentrations of sulfur dioxide (SO2) and ozone (O3) are observed in SAG (Fig. S9). The higher concentrations of SO2 and O3 in SAG, which is a smaller urban center, are related to the presence of several large industrial complexes (*18, 27*). Coinertia analyses (*28*), revealed covariation between 57 clinical endophenotypes (Table S4), environmental exposures (Fig. S12), and expression levels of DEGs and RDEGs (Fig. S10). Consistent with documented effects of air pollution on cardiac and respiratory traits (*29*, *30*), we found that arterial stiffness measures, asthma and stroke prevalence, monocytes counts, low-density lipoprotein (LDL), respiratory function (FEV1), as well as liver enzyme levels (Alanine aminotransferase level (ALT), Aspartate aminotransferase level (AST), Gamma-glutamyl transferase (GGT)) show the strongest associations with SO2 and O3 ambient levels (Fig. S10).

**Figure 3:**
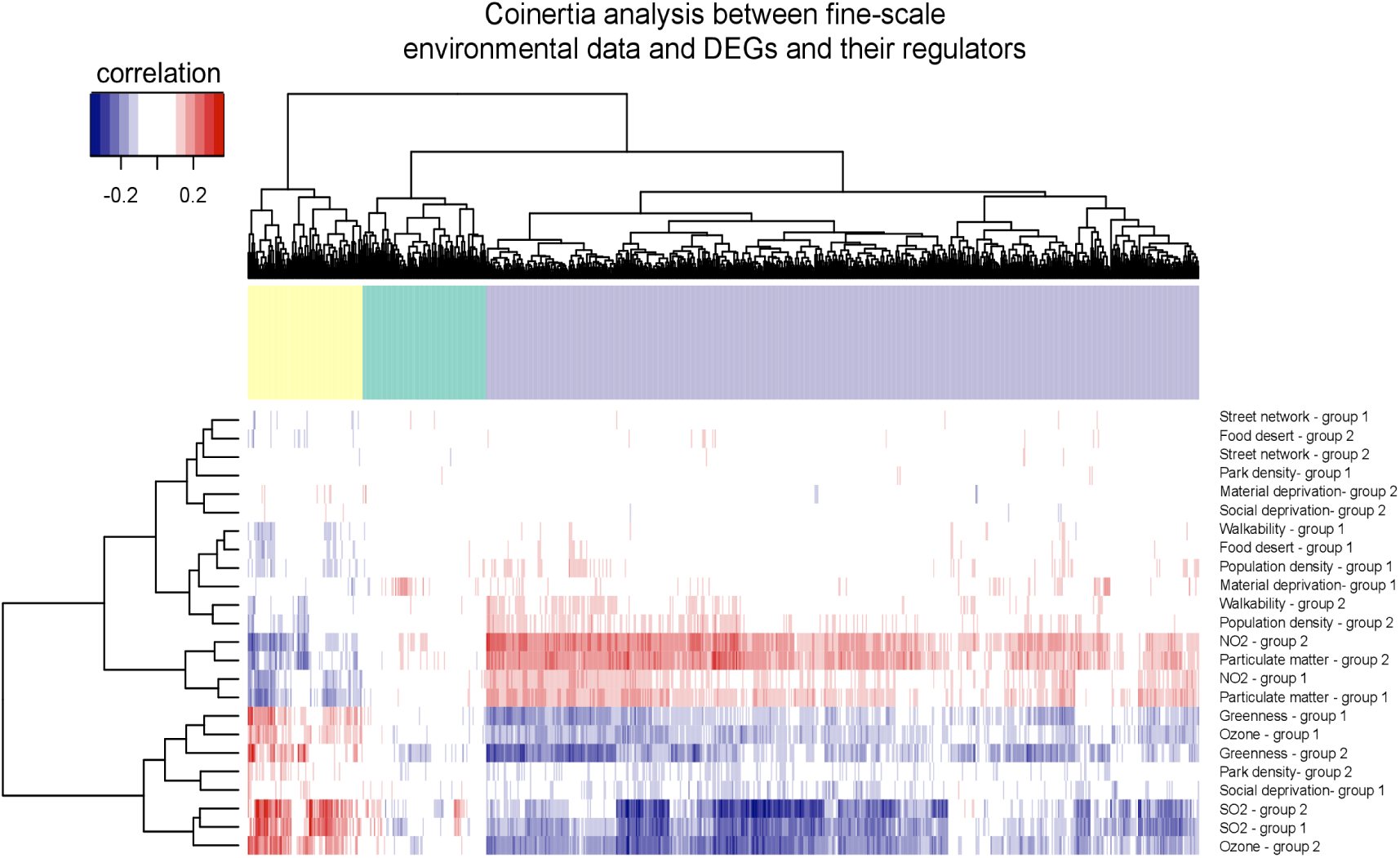
Differentially expressed genes are associated with local ambient air pollution. Coinertia (CoIA) analysis between gene expression (columns) and fine-scale environmental variables (rows). CoIA analysis were performed on genes that were significantly differentially expressed among regions and the regulators of those genes (RDEG). CoIAs were computed between DEGs profiles and fine-scale environmental data (Fig. S11). We performed two sets (Group 1 and Group 2, each composed of a random draw of half the cohort) of CoIAs: each set included 10000x resampling of 200 individuals (without replacement), and the CoIAs were performed between environment and gene expression for each of the 10000 iterations. Figure SS11 depicts the resampling scheme. The above heatmap represents, for each Group 1 or Group 2,the median of each environment-gene associations from the cross-tabulated values distribution. Associations from Group 1 and Group 2 largely cluster together, indicating a strong signal of the association between fine-scale air pollution levels and gene expression. A permutation test (n=10000 steps) indicates the that the correlations between the matrices are significant (p=0.00089 and p=9.9e-05 for group 1 and 2 respectively)

We increased our resolution for pollution exposures by using daily exposure to SO2 pollution averaged over a 14-day window preceding each individual sampling day (Fig. S14). The large temporal fluctuations in SO2 ambient concentrations over time scales of a few weeks allowed us to include individuals from SAG that were exposed to low levels of SO2 (despite SAG having high annual averages), and MTL individuals exposed to high levels of SO2 (despite MTL having lower annual averages), or vice-versa. In that way, we reveal the effects of the local environment specifically due to recent SO2 exposure and uncorrelated to broad regional sampling. Using a robust resampling design to balance the number of individuals in each category (See Supplementary materials), we identified 170 DEGs between high- and low-exposure individuals, which were also found as DEGs between regions (Fig. 2A), supporting their association with air pollution (Fig. S10). Furthermore, while multivariate models show that gene expression variation for those 170 genes is significantly associated with SO2, they do not show an association with smoking or socio-demographic status, or with most built environment characteristics (Table S6). We performed a sensitivity analysis using MTL-only samples, thereby removing the potential influences of geographic region and regional ancestry. We replicated these associations with pollution, and the lack of thereof for smoking and socio-demographic status (Table S6). Those 170 DEGs are again enriched in oxygen-transport activities, and in several pathways involved in leukocyte migration during chronic inflammation, including CXCR chemokine activity and G-protein coupled receptors (Table S5). Circulating blood leukocytes migrate to sites of tissue injury by responding to proinflammatory cues (*31*), and are known to migrate through the blood flow to lung epithelial cells during inflammatory response (*31*).

Using the endophenotypes identified with CoIA as being associated with air pollution (Fig. S10), multivariate models also show that gene expression of the 170 DEGs is significantly and strongly associated with several of these traits (FEV1, lung disease, liver enzymes, arterial stiffness), across Quebec, and also within Montreal. Furthermore, when the effects of these significant endophenotypes are regressed out from gene expression, SO2 remains significantly associated with gene expression (Table S6), suggesting that exposure itself is primarily modulating gene expression, and not underlying health status. Most of the endophenotypes that we found to be associated with DEG expression, such as pulmonary function and arterial stiffness, are consistently reported as influenced by air pollution (*32*–*35*). GGT, which we found to be associated with the expression of genes enriched in blood coagulation and platelet regulation (Fig. S12), has been found in atherosclerotic plaques (*36*) and is elevated following pollution exposure (*37*, *38*), and is predictive in a dose-dependent manner of cardiovascular risk (*39*). Collectively, these results reveal associations between environmental pollutants, endophenotypic traits, as well as transcript levels, and that the type and direction of associations are consistent with detrimental effects of air pollution, or a correlated variable, on health status.

Environmental factors not only directly affect phenotypic variation, but can also modulate associations between segregating genetic variants and phenotypes (*1*, *40, 41*). To discover gene-by-environment interactions in both FCs and Europeans, we identified eQTLs for which the effect size is modulated by exposure with one of four ambient air pollutants (env-eQTLs): PM2.5, NO2, O3, and SO2. First, we identified canonical eQTLs using 5,313,384 genotypes (genotyped or imputed - see Supplemental material) and show a high replication for proximal canonical eQTLs (cis-eQTLs) with previously discovered cis-eQTLs (Table S7).

We identified 34 environmentally responsive genes (env-eQTLs eGenes) in FCs, that are modulated by pollution exposure, and replicated in Europeans. We identified these environmentally responsive genes by using gene-specific bonferonni corrected *p*-values (Fig. 4A, Table S8). The expression of those 34 eGenes are modulated by cis-SNPs, and exhibit significant interactions with a pollutant level. Significance was assessed with gene-specific bonferonni corrected p-values and associations were not driven by outlier individuals (Fig. 4, Fig. S15, Table S8). The correlations of env-eQTLs effect sizes between FCs and Europeans are positive, large, and significant (Fig. S17). Among our most significant env-eQTLs (Fig. S16), we identified an interaction with NO2 and the SNP-gene pair rs10814466-PAX5 (Fig. 4). PAX5 gene is a transcription factor expressed in leukocytes and has been associated with asthma in a population cohort from Saguenay-Lac-Saint-Jean (*42*). Together, these results document a number of environmentally responsive loci for which individual genetic variation modulates expression levels and may be associated with clinical conditions, such as asthma, which have different prevalences across Quebec (Fig. S10) (*43*).

**Figure 4:**
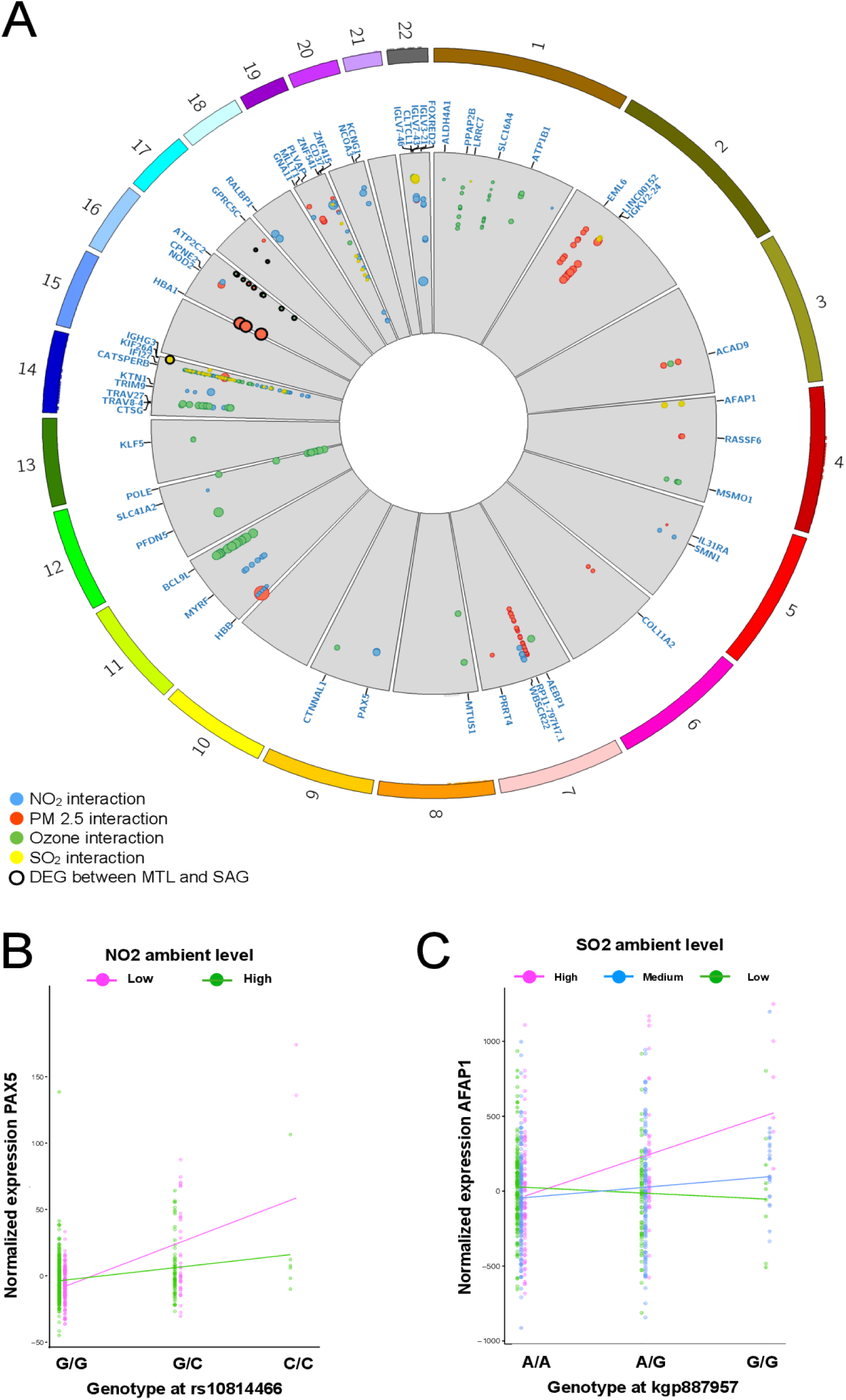
Genome wide env-eQTLs map in CARTaGENE. (A) Circular manhattan plot displaying all significant cis-eQTLs for which an interaction between the genotype and an environmental air pollutant was identified (env-eQTLs). eGenes shown here replicate between analyses performed with either FCs or European individuals. Each point represent a SNP-gene pair that was significant, its position relative the center reflects the −log10(p-value) (higher closer to the center), the colour represents eQTLS with significant pollution interactions, and the size of the dot reflect the log10 effect size of the eQTL. Bold dots represent genes that were also found to be differentially expressed between MTL and SAG. Examples of interactions of the genotype and the environment on expression level of (B) PAX5, and (C) AFAP1.

We reveal a signature suggesting that rare genetic variation has a disproportionate effect on phenotypic variation, and in particular on rare variant eQTLs susceptible to modification through environmental stimuli. First, we document that large effect sizes on transcript abundances in env-eQTLs are predominantly mediated by rare eSNPs (Fig. S18). This pattern is consistent with natural selection acting to stabilize gene expression (*44–46*), and that rare variants mediate larger changes in phenotypic trait variance particularly when influenced by environmental perturbations (*46*, *47*). Second, we performed a CoIA analysis between endophenotypes showing differences across environments (Fig. S10) and env-eQTL eSNPs genotypes. Rare variants (MAF < 0.1) are overrepresented for strong associations between eSNPs genotypes and endophenotypic traits (Fig SS19), suggesting that rare variants responding to an environmental stimuli may mediate larger phenotypic changes than common variants.

Lastly, we find evidence that suggests that personal disease risk can be modulated by rare genetic variants influencing expression levels of genes implicated in chronic diseases, together with environmental exposures. For example, we identified that PAX5 and CTSG, two genes that have been associated with asthma etiology (*42*, *48*), exhibit gene-by-air pollution interactions. Individuals carrying homozygous derived genotypes show larger changes in expression level in response to a high pollution exposure than when exposed to a lower level (Fig. S15). Such personal genetic sensitivity profiles to environmental exposures may help develop individualized care for the prevention and treatment of chronic diseases (*49*).

Our findings illustrate that the impact of the geographic region of residence on the blood transcriptome overrides that of ancestry. Moreover, ambient air pollution exposures are likely contributing to this regional effect in Quebec and may explain the differences in some clinical traits among regions such as asthma prevalence. Fortunately, in Quebec and in many parts of the developed world, air quality has improved since the 1980s (*27*, *50*). However, there has been a sharp increase in anthropogenic pollution levels in many parts of Asia caused by the rapid industrialization and increased use of fossil fuel energies. In the context of global climate change, air pollution and hazardous air quality events are predicted to become more frequent and cause additional morbidity and mortality (*19*). More broadly, our work shows how environmental exposures modulate gene expression directly and can drastically affect clinically relevant phenotypes in humans.

## STAR Methods

### CONTACT FOR REAGENT AND RESOURCE SHARING

Further information and requests for reagents may be directed to the Biobank CARTaGENE which regulates the access to the data and biological materials.

### EXPERIMENTAL MODEL AND SUBJECT DETAILS

#### Study population

The study protocol was approved by the Ethical Review Board Committee of Sainte-Justine Research Center and all participants provided informed consent. CARTaGENE biobank comprises of more than 40,000 participants aged between 40-60 years old, recruited at random among three urban centers in the province of Quebec. CARTaGENE is a regional cohort within the Canadian Partnership for Tomorrow Project, including over 315,000 participants, with various measures obtained from blood parameters, biological function, disease history, lifestyle, and environmental factors (*15*).

#### Sample selection

For set 1, we selected 708 individuals from the CARTaGENE’s biobank samples with available Tempus Blood RNA Tubes (ThermoFisher Scientific) and Framingham risk scores, ensuring an equal representation of ages and gender. Two-hundred-and-ninety-two additional samples were subsequently selected in CARTaGENE’s biobank based on their RNA and complete arterial stiffness (AIx) measures availability. These samples were selected for having high AIx values as well as average AIx values to complete the first set of samples in the intention of achieving a range of arterial stiffness values across the study cohort.

### METHOD DETAILS

#### Genotyping, ethnicity, and regional origin of French-Canadians

928 samples with RNA-Seq profiles that passed quality control (QC) thresholds were genotyped on the Illumina Omni2.5 array to obtain high density SNP genotyping data. A total of 1,213,103 SNP were retained after filtering and QC (Hardy-Weinberg p-value > 0.001, MAF > 5% and percent of missing data < 1%).

#### RNA sequencing

Whole blood samples were collected from participants in 2010. Total RNA was isolated using the Tempus Spin RNA isolation kit (ThermoFisher Scientific) and a globin mRNA-depletion was performed using the GLOBINclear-Human kit (ThermoFisher Scientific). The quality and integrity of the RNA samples were verified using an Agilent Bioanalyzer 2100 and all samples had an RNA Integrity Number (RIN) > 7.5. A RIN above 7.5 is indicative of high quality RNA in the sample and for which RNA degradation is minimal, indicating optimal transport and preservation conditions. Our RIN threshold is more stringent than other large scale consortium studying gene expression in tissues (*44, 51*). TruSeq RNA Sample Prep kit v2 (Illumina) was used to construct paired-end RNA-Seq libraries with 500ng of globin-depleted total RNA. Recommended Illumina protocols were followed for quantification and quality control of RNASeq libraries prior to sequencing. Paired-end RNA sequencing was performed on a HiSeq 2000 platform at the Genome Quebec Innovation Center (Montreal, Canada). Sequencing was performed for set 1 (708 samples) using 3 samples per lane, and for set 2 (292 samples) using 6 samples per lane.

Reads were trimmed for adapters and bad quality bases first using Trim Galore and were then assembled to a reference genome (hg19, European Hapmap (CEU) Major Allele release) using STAR (v2.3.1z15) (52) using the 2-pass protocol, as recommended by the Broad Institute. The 2-pass protocol consists in two consecutive mappings steps having the same set of parameters with only the reference that is optimized in the second mapping procedure. The first mapping is done using the reference gene definition coming from ENSEMBL (release 75). Then, using the splicing junction database files formed by the first pass mapping step for all the samples combined together and the same gene definition file, a second reference is indexed and optimized and is used for the second mapping step. The number of mismatches allowed across pair is five and a soft-clipping step that optimizes alignment scores is also done automatically by STAR. The PCR duplicates were conserved as it was shown that quantification of highly expressed genes were disproportionately affected by pcr duplicates removal (53). Only properly paired reads were kept (using samtools (*54*)) for the analysis, according to STAR parameters. After these steps, HTseq (v.0.6.1p1) (*55*) was launched separately on each alignment file using the same gene reference file that was used for the alignments.

All analyses downstream were conducted using R 3.1.2 and R 3.2.2 and Bioconductor R packages.

### QUANTIFICATION AND STATISTICAL ANALYSIS

#### Fine scale population genetic structure within French-Canadian population

To unveil finer scale patterns of population structure, i.e. differences between individuals with European ancestry versus individuals having a French Canadian ancestry, we also used ChromoPainter (v0.04) (*26*), a haplotype-based method powerful enough to detect fine-scale genetic structure. Original genotyping data was used apart from singletons, yielding to 1,908,336 SNPs. Singletons were removed as they are non-informative for phasing and contribute to computation burden for the step of haplotypes sharing inference performed with ChromoPainter. Genotypic data was phased with SHAPEIT (v2.r644) (*56*) using the HapMap genetic maps. Coancestry matrices were obtained from ChromoPainter with parameters estimation step done with 10 iterations on four chromosomes only. ChromoPainter method performs a reconstruction of every individual genome using chunks of DNA donated by the other individuals and report matrices of the number and length of those chunks. We used the chunk count matrix to (1) run FineSTRUCTURE algorithm to build a tree (as recommended for large dataset, we performed 10,000,000 burn-in and runtime MCMC iterations) (Fig. S1D) and to (2) perform a PCA (Fig. 1A, Fig. S1C). Regional ancestry for each FC was determined based on the three clusters obtained from the fineSTRUCTURE tree, (Fig. S1D, Fig. 1B).

In agreement with Quebec settlement history, previous studies of the Quebec population (*21, 25*, and the fineSTRUCTURE tree, a PCA of French-Canadian (FC) individuals reveals groupings of sub-populations of individuals that follow a North-South cline (Fig 1B and C). The founding event from French settlers followed by the subsequent colonization of remote regions has led to population differentiation among regions in Quebec (*21, 25*). By further restricting the group of individuals to be analyzed to only FC (n=726) and considering their region of residence (either Quebec City, Montreal and Saguenay) a PCA on the chunk count matrix reveals three groups corresponding to region of residence, with the Montreal and Quebec groups overlapping to a greater extent, in line with their greater geographic proximity (Fig. 1B and C). Those three groups were also recovered by the fineSTRUCTURE tree (Fig. S1D). Considering all SNPs and the whole haplotypic structure is the key in seeing differences for those two metropolitan regions that have low differentiation. We further identified several participants with a regional ancestry discordant with their region of residence: an indication of recent internal migration of these participants across Quebec regions (Table S2).

#### Imputation

To increase the power for the association study with gene expression levels, variant imputation was conducted on 968 individuals for which the genotyping was available from the Illumina Omni2.5 array. We pre-phased the genotypes with SHAPEIT (v2.r64410) (*56*) using the default parameters, on both the autosomes and the chromosome X. We filtered variants for MAF > 1% and Hardy-Weinberg p-value > 0.0001 and passed the haplotypes to IMPUTE2 (v2.2.2) (*57*) to perform the imputation using the 1000 Genomes Phase I integrated haplotypes (Dec 2013). We used the parameters Ne = 11418 and call_thresh = 0.9. We removed variants with a call rate less than 90%, MAF > 1% and Hardy-Weinberg p-value > 0.0001. A total of 9,157,622 variants passed the filters. Of these, 8,877,297 variants were found on the autosomes and included 779,579 insertion-deletion polymorphisms (indels) (8.78%) and 8,097,718 SNPs (91.22%). 280,325 variants were found on the chromosome X, which included 28,504 indels (10,16%) and 251,821 SNPs (89.84%).

To determine the ancestry of each individual from genotyping data, we carried out a principal component analysis (PCA) with SNPs pruned for LD (pairwise r2 > 0.2 and 50 SNPs window shifting every five SNPs) (Fig. S1A), yielding 146,689 SNPs. The continental ancestry (African / European / Asian / Canadian / American / Middle-Eastern) of each individual was determined based on the PCA plot (Fig. S1A) and verified as to whether it corresponds to self-reported ancestry based on the country of origin of four grand-parents. If the country of origin of three out of four grand-parents and the PCA continental grouping were concordant, the individual was assigned to a continental origin.

#### RNA-sequencing filtering

Genes with counts-per-million below 0.5 in more than half of the cohort (505 individuals) were removed from the analysis for a total of 15632 genes retained for all downstream analyses. Individuals that showed obvious outlier after visual inspection of principal component plots were removed (3 individuals). Pre-processing and normalization of the raw gene counts matrix for the remaining 708 (or 289 in set 2) samples was done using Bioconductor’s library EDASeq (v 2.0.0). Gene counts were corrected for GC-content bias and gene length using full-quantile normalization between feature strata as described in (*58*). The resulting expression matrix was quantile normalized across samples to account for library size variation among samples.

#### Exploration of variables contributing to transcriptomic variation

The deep phenotyping of the CARTaGENE cohort allow for a thorough exploration of the biological and environmental factors that may influence genome-wide gene expression patterns. We applied the low expression filters and normalized the counts from set 1 according to the pipeline described above (EDASeq normalization for gene length, GC content and library size). As most statistical procedures assume a normal distribution to the underlying data, we transformed the normalized counts from set 1 to a gaussian distribution using a log2cpm transformation using edgeR. We summarize the gene expression levels by performing a PCA on the normalized expression matrix (ePCA). To identify variables that contribute to genome-wide gene expression variation, we performed a stepwise regression (stepwise search from both directions) on ePC1 and ePC2. Results of the stepwise regression are given in Table S1, as well as the results from the replication analyses using set 2. We included the following low level endophenotypes in the stepwise procedure: set, region of residence, cell counts (lymphocytes, neutrophils, monocytes), arterial stiffness, age, and sex.

#### Sampling site effect within region

The RNA extractions and library preparation were performed for all individuals in the same laboratory to reduce technical bias. However, the participants were sampled across four different sampling sites inevitably situated within the geographical regions where the participants lived. Our experimental design was built in such a way that sequencing run was not correlated with region of residence (Fig. S3A). To evaluate whether the sampling site has any effect on the RNA-Seq quantification data, we performed extensive analyses of the two sampling sites situated within Quebec city: St-Sacrement (STS, n=136) and Enfant-Jesus (EF, n=129). QUE individuals expression profiles from the combined dataset show that individuals from STS and EF form a single cluster on a MDS plot (Fig. S3B). We show that the set (discovery or repli-cation) has a greater impact on the expression profiles than the sampling site (Fig. S3C). Furthermore, a variance component analysis (PVCA) was performed on the QUE individuals only and including sampling site as an explanatory variable shows that the sampling site explains less than 5% of the variance within QUE region, while set explains 15%, age 5% and gender, 2.5% (Fig. S3C). In comparison, in FCs or Europeans, region of residence accounts for 15% of variance in gene expression.

#### Correction for technical and biological unwanted variation

RNA-Seq data generation, and expression data in general, are prone to technical biases which in some cases can mimic, or be confounded with biological variation. The appropriate normalization pipeline in an RNA-Seq experiment will depend on the experimental design and the hypothesis being tested. Local sequence context can bias the uniformity of read counts along the genome, and sophisticated normalization pipeline may be necessary when comparing expression levels across genes (*59*). Most experimental designs of RNA-Seq studies, like the one presented here, compares different groups of individuals to each other, therefore the normalization pipeline should rather focus on removing unwanted variation across individuals. We removed potential effects of hidden covariates potentially affecting expression levels using surrogate variable analysis (SVA) (*60*) and probabilistic estimation of expression residual (PEER) correction (*61*). We show a comparison of SVA and PEER on removing unwanted variation in the gene expression data in Table 1. We used the SVA correction, retaining 1 surrogate variable, for the differential expression analyses. We performed the same stepwise regression approach as previously, but on the SVA and the PEER corrected expression level matrices and show that we retained the variation associated with region, but removed any effects of cell counts and arterial stiffness that was present in the uncorrected expression levels (Table S1). The corrections do not fully compensate for the effect of the set (technical), we therefore include this covari-ate in all subsequent analyses. PEER approach has been shown to remove variation associated with biological and technical factors and also increase the power to identify eQTLs (*53*, *62*). The choice of the number of PEER factors we removed (k=15) is described below in the eQTL section. We specifically retained variation associated with region. We show that a number of biological and technical factors do indeed correlate with each of the PEER factors (Fig. S4).

#### Differential expression analysis between regions and regional ancestries

Because of the large proportion of the variance in gene expression explained by region of residence revealed by the stepwise regression, we then identified genes that are differentially expressed between pairwise comparisons between the FC-locals from the 3 regions (Montreal, Quebec and Saguenay). Using edgeR (*63*), we performed a differential gene expression analysis using the 15632 genes that passed the QC filters established above. We performed the differential expression modeling using the following statistical model:

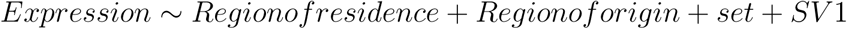

The significance level of the test was estimated as a gene p-value below the bonferroni-corrected threshold at 3.20 x 10^−6^ (0.05/15 632). We then performed the same stepwise regression approach as previously, but on both the SVA and PEER corrected expression levels and show that we retained the variation associated with region, but removed any effects of cell counts and arterial stiffness that was present in the uncorrected expression levels (Table S1). Both the SVA and PEER correction does not fully compensate for the effect of the set (technical), we therefore include this covariate in all subsequent analyses requiring covariates. We performed a power analysis of our ability to detect differentially expressed genes with smaller samples sizes. Several of our comparisons of local migrants or continental migrants with FC-locals involve smaller number of individuals (Table S2). We therefore assessed our ability to detect DEGs by performing differential expression analyses between groups for which we found large number of DEGs, but using a smaller subset of random individuals (without replacement) of each of these groups. We randomly selected 15 Mtl-locals and 15 Sag-locals, and performed the differential expression analysis using the same model as above. We also performed the analysis using 50 Mtl-locals and 50 Sag-locals. In each case, we could identify DEGs which largely overlap with the DEGs detected in comparisons using all individuals (Fig. S6A to C). We observe that with an increasing number of individuals, our power to detect DEGs increases and that the identity of the differentially expressed genes detected in each of these comparisons largely overlap (Fig. S6A to C).

#### Regional environmental effects on gene expression

We take advantage of the presence of individuals from different regional and continental origins in our cohort to disentangle further the effects of the genetic background and environmental influences on genome-wide gene expression. We first selected individuals of either French-Canadian and European continental ethnicity (Fig. 1A, Fig. S1A). A total of 798 individuals including 136 Europeans and 662 FC were selected for downstream analyses. We stratified the individuals according to their continental origin (FC vs Europeans), and further stratified the FCs into their assigned genetic ancestry (MTL, QUE, SAG) obtained from the fineSTRUCTURE analysis (Fig. 1B, Fig. S1D). We then determined their region of residence (MTL, QUE, SAG) for a total of 12 ancestry-residence groups: we identified individuals for which their origin (Continental or regional) is discordant with the region they reside, which we refer to as continental-migrants and regional-migrants respectively (Table S2). We also identified FC individuals for which their regional origin is concordant to the region they reside, which we refer to as FC-locals (Mtl-FC-locals, Que-FC-locals and Sag-FC-locals). We performed the differential gene expression analysis pipeline as described above for different pairs of continental-migrants, regional-migrants, and FC-locals to disentangle the effects of the genetic background and the regional environment on genome-wide expression (Fig. 2). We selected 6649 genes that show differential expression (p-value < 3.20 x 10^−6^) in the comparison between Mtl-FC-locals and Sag-FC-locals. Using the 12 origin-living groups and the 6649 genes, we performed an unsupervised clustering and visualized the groupings using a heatmap (Fig SS5).

#### Gene enrichment and Reactome analyses

Gene enrichment analyses were performed using the topGO package in R, with a classic fisher test. Differentially expressed genes between MTL-locals and SAG-locals were compared against the 15632 genes expressed in the CARTaGENE cohort that were retained after QC filters (background). Reactome enrichment analyses were conducted with R the package reactomePA, and here again, the background set of gene was defined as the 15632 genes expressed in blood that pass our filters (Fig. S7A and B).

#### Fine-scale environmental data

We obtained air quality measures in the year of sampling (2010) from either land-based stations (SO2, ozone) or national LUR models estimates (PM 2.5 and NO2) incorporating information from land use data and satellite remote sensing (*50*, *64*–*66*). Built environment variables (street network, population density, food deserts, greenness, walkability) and social and material deprivation indicators were accessed through the Quebec government data portal (https://www.inspq.qc.ca/environnement-bati)

Environmental data was available at the 3-digit postal code district level (i.e Forward Sortation Area, FSA), or was reformatted to this geographic scale. Postal code districts in Canada are small geographic areas which assist in delivering mail. Postal codes are a series of 6-digits that identify a small geographic area in a municipality, usually grouping just a few houses together or a small neighborhood. 3-level digits are larger areas that include several houses, a small neighborhood, or a small village. The population of FSAs in Canada range from a few hundreds to tens of thousands of individuals. 3-digit postal code districts can be of different areas, and are smaller in densely populated areas, and larger in areas of low population density. Maps in Fig 1C and D, and Fig SS9B to E depict 3-digit postal code districts as thin grey lines areas, and each district is colored with the mean value of interest in each map. Each individual in the CARTaGENE cohort has a 3-digit postal code district associated to it, referring to the location of its primary residence. We assigned fine-scale environmental measures to each individual based on its 3-digits postal code.

#### Coinertia analyses (ColA)

Coinertia analysis (CoIA) (*28*, *67*) is a multivariate statistical part of the large family of ordination methods, such as principal component analysis (PCA), redundancy analysis (RDA), or canonical correlation analysis (CCA). CoIA is a general approach and existing methods such as the ones mentioned above appear as special cases of it (28). These methods have been widely used in ecological research, including CoIA which has been more recently developed. Collectively, these methods allow for detecting an underlying data structure between two data tables. CoIA uses a combination of PCA and multivariate linear regressions to detect linear combinations of variables from one data table that explain the variance in the second data table. CoIA is more flexible than RDA or CCA, and overcomes their limitations by allowing for more variables than the number of samples to be tested (*28, 67*), which is generally the case in genome-wide scale analyses (i.e. more genes than individuals).

We first used coinertia analysis to reveal the common structure between DEGs plus the genes that regulate them (Fig.2) and the fine-scale environmental data (Fig. S8). We produced two separate principal component analyses (PCAs) based on continuous encoded matrices of both environmental and gene expression levels (normalized for library size and sequencing freeze). We conserved components for each PCA to explain 80% of the variance in the data. We imputed missing data (in the fine-scale environmental data, there were no missing data in the gene expression matrix) using the function *imputePCA* from the R package missMDA. In the CoIA, the separate PCAs were rotated to a comparable alignment and normalized co-inertial loadings were obtained for each environmental variables or gene expression. Relationships between the two matrices were assessed by comparing the CoIA estimated from the real data set with the CoIA distribution estimated after bootstrapping. Two sets of 500 of CoIAs were computed independently between gene expression and fine-scale environmental data. Figure SS11 depicts the resampling scheme. For each Group 1 or Group 2 (n=497 for each) a total of 10000x resampling of 200 individuals (without replacement) were performed. We performed a CoIA for each resampling step. We report the median value of the distribution of each environment-gene expression pair cross-tabulated values for each group. Gene enrichment were performed using gProfiler (*68*), and using the 15632 expressed genes that passed our filters in whole blood as the background gene set (Fig. 3). We evaluated the significance of the correlations between the matrices using a permutation test (RV-test) with 10000 steps form the R package ade4.

To identify clinically relevant endophenotypes that are associated with fine-scale environmental data, we performed a CoIA between 57 clinically relevant endophenotypes (Fig. 4) and fine-scale environmental data (Fig. S8). The 57 clinically relevant endophenotypes were selected to encompass physical measures (BMI, height, age, sex), most systems relevant to the human health (cardiovascular system, pulmonary functions, hepatic system, renal system, disease history, vision, immune system) and lifestyle measures (smoking status, alcohol consumption, nutrition, physical activity). All biochemical endophenotypes were measured in a single central laboratory. We resampled 10000 times 493 individuals from the cohort, and performed CoIA at each step between endophenotypes and fine-scale environmental variables. We report the median value of the distribution of each environment-endophenotype pair cross-tabulated values (Fig. S10).

To reveal possible associations between expression levels and endophenotypes, we then performed CoIAs with a similar resampling scheme (SS11) between 12 selected endophenotypes that were the most strongly associated with air pollutants (Stroke, Arterial stiffness measures, spirometry measures, Asthma, monocyte counts, LDL, AST, ALT, GGT) and differentially expressed genes (DGEs and RDEGs) (Fig. S12).

To find associations between endophenotypes and env-eQTL eSNP genotypes, we also performed CoIA between selected endophenotypes that were the most strongly associated with air pollutants (Stroke, Arterial stiffness measures, spirometry measures, Asthma, monocyte counts, LDL, AST, ALT, GGT), and env-eQTL eSNPs. We resampled 420 individuals 10000 times and performed CoIAs between endophenotypes and env-eQTLs eSNPs discovered in FCs and calculated the median for each eSNP-endophentoype pair. In this case, we resampled a smaller number of individual because the total number of individual with genotyping data dos not include all individuals (n=928). To assess the possibility that rare SNPs are associated with larger endophenotypic changes when under the influence of an environmental stimuli, we calculated the odd ratios of rare eSNPs (MAF < 0.1) showing larger cross-tabulated values in the CoIA (Fig. S19). Large odds ratios are indicative of an enrichment of rare SNPs for strongest associations.

#### Daily SO2 data: exposure windows

To increase our resolution in air pollution exposures, we used daily SO2 ambient levels measured in each mail sortation area. We calculated the average exposure during the two weeks preceding the blood draw for each participant. This way, we reduce the effect of random fluctuations due to technical artifacts or short-term meteorological anomalies that may affect measurements. Also, changes in gene expression and biomarkers in blood following a pollution exposure has been documented as a relatively fast phenomenon, occurring after just a few days of exposure (*33*). We then categorized the participants into high exposure (over 2) or low exposure (below or equal 2) categories.

#### Daily SO2 data: DEG between high and low exposure individuals

To find differentially expressed genes between high and low exposure individuals, we used the same approach as described above for identifying DEG between regions, with the following modifications: given the unbalanced number of individuals in each category (108 high exposure vs 800 low exposure) of exposure, we resampled 100 times 108 individuals from the low exposure category and performed the DEG pipeline. We performed the SVA while retaining variation associated with SO2 exposure. We combined the results of DEG in a list of 468 DEG, and from these candidates, 170 genes were also identified as DE between regions (2A). Those strong 170 candidates were used for enrichment and multivariate models.

#### Daily SO2 data: multivariate models

In an effort to characterize the effects of confounding variables on pollution exposure, we performed multivariate models on gene expression levels. First, similar as in the DEG, we performed a SVA to remove unwanted variation of technical or unknown biological variables while retaining the variation around SO2 exposure. We then built multivariate models using the SO2, O3, and PM2.5 14-day exposures, as well as the remaining 9 non-pollution environmental exposures (Fig. S9), as well as smoking status (Fig. 4). Smoking status may indeed cause similar changes in endophenotypes as pollution exposure. We then selected the endophenotypes revealed by the CoIA as being the most associated with region and pollution exposure (Lung disease, Asthma, Stroke, monocyre counts, liver enzymes (AST, ALT, GGT), Arterial stiffness, spirometry tests, and lymphocyte counts, Fig. 4), and tested whether any of these would explain variation in the 170 candidates. Furthermore, after having identified the health endophenotypes that are associated with gene expression in Montreal and in the whole data set (FEV1, liver enzymes, lung diseases, and arterial stiffness, see Table S6), we regressed out their effect from the expression of the 170 candidate genes, and run the multivariate models to test for the effects of environmental variables. We found that daily SO2 exposure still explain significant variation in gene expression after removing the effect of health endophenotypes, in both MTL and the whole data set, suggesting that SO2 exposure itself changes expression, and not the health endophenotypes associated with it.

#### Canonical QTLs and env-QTLs

As discussed above, we corrected expression levels with PEER for the eQTL analyses. To select the appropriate number of PEER factors that maximizes our power for cis-eQTL detection, we run the PEER correction sequentially removing 5, 10, 15, 20, 25, 30, 35 and 40 factors and calculated the expression residuals from the inferred PEER factors excluding the region covariate to preserve the effect of region on expression. To decide of the number of factors to remove, we follow (*62*). We applied the interaction eQTL pipeline, using the non-imputed genotyping data to reduce computational burden. We found that removing k=15 PEER factors maximizes the number of eQTLs detected. To verify the effect of the PEER correction in our dataset, we (1) show that PEER factors indeed correlate with many variables that were measured in the cohort, both technical and biological (Fig. S4) and (2) performed the stepwise regression using the PEER corrected data for which the effect of region has been retained. Indeed, after correction, only region and sampling set remained significant (all effects of cell counts and arterial stiffness on gene expression variation have been effectively removed by PEER) (Table S1). We therefore use PEER corrected expression data for all subsequent eQTL analyses.

For the detection of eQTLs (Table S7), we selected participants of FC ancestry for which we had gene expression data and that passed QC filters for the gene expression and imputed genotyping data (participants n=689, HW p-value > 0.001, MAF > 0.05 in each region and percent of missing data < 1%, for a total of ∼ 5M SNPs). We used the 15632 genes that passed our QC filters as quantitative phenotypes. To detect linear canonical eQTLs, we tested the following model in FCs and Europeans:

Linear eqtls:

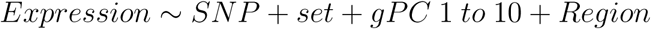

We defined a cis-eQTL as a SNP falling within 1Mb up- or downstream of a gene and influencing its expression. cis-eQTLs were mapped using the R package MatrixEQTL using the linear (for canonical eQTLs) or cross-linear (for air pollution interaction) models. We used bonferroni-adjusted thresholds for significance, which were calculated as follow. Depending on the dataset used (FC, Europeans), the thresholds changed because of the different number of snps in each gene in each dataset.

cis-eQTLs: 0.05/(15632* N snps in gene filtered for LD)

We then added an interaction term to detect gene-by-environment interactions, with four ambient air pollutant levels that we categorized (see below): PM 2.5 (particulate matter 2.5), NO2, ozone, and SO2, using the following models:

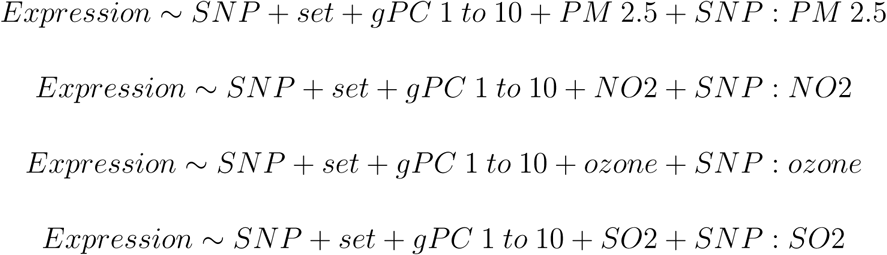

The pollution ambient concentrations were binned into categories for the env-eQTL modeling as follow:

|  | Low | Medium | High |
| --- | --- | --- | --- |
| PM 2.5 | equal or below 8 | - | above 8 |
| NO2 | equal or below 11 | - | above 11 |
| Ozone | equal or below 23.5 | - | above 23.5 |
| SO2 | equal or below 1 | Between 1 and 2.5 | above 2.5 |

#### Reporting env-eQTL significant interactions

We report in Table S8 the number of significant env-eQTL eGenes for interactions with each pollutant. We obtain relatively large numbers of significant eGenes when we tested independently in FCs and in Europeans (Fig. S15). To increase confidence in the reported interactions, we identified the eGenes that replicate in both FCs and Europeans, for each pollutant, reducing significantly the number of interactions identified (Fig. 4), and strengthening confidence in those filtered interactions. We further refined our poll of significant interactions by graphing the genotypes, the expression level, and the environmental exposure for each individual (examples shown in Fig. S16), and visually assessing the relationships (and identifying possible associations driven by few outliers, or for which the effect size was very small). This last set of interactions are the ones for which we have the highest confidence in (n=34).

### DATA AND SOFTWARE AVAILABILITY

Genotyping, expression, health phenotypes, and exposure data used in this study are available from the CARTaGENE Biobank upon request. The built environment data is publicly available from the Quebec government data portal. The air pollution data is available upon request to Air health Effects division, Government of Canada.

## Author contributions

Conceptualization: M-J.F., P.A., A.J.H, Y.I. Experimental procedures for sequencing and geno- typing: E.G., Y.I. Data preparation and quality control: V.B., J-C.G., M.J., A.S. Bioinformatic and statistical analyses: M-J.F., F.C.L., V.B., J-C.G, H-G. Manuscript writing and revising: M-J.F, F.C.L, K.S, P.A.

## Acknowledgments

We thank the Awadalla lab for comments on the manuscript, as well as Paul C. Boutros and Veronica Y. Sabelnykova from OICR. We acknowledge financial support from Fonds de la recherche en sante du Quebec (FRSQ), Genome Quebec, Fonds quebecois de la recherche sur la nature et les technologies (FQRNT), Canadian Foundation of Innovation, Ontario Ministry of Research and Innovation Principal Investigator Award and a Canadian Institute of Health Research award (#EC3-144623) to PA. M-JF is a CIHR Neuroinflammation Postdoctoral Fellow. AH is an FRSQ Postdoctoral Fellow and currently holds an eMedLab Career Development Fellowship as part of the Medical Bioinformatics Initiative funded by the Medical Research Council, UK (grant number MR/L016311/1). FCL is a FRSQ Postdoctoral Fellow. Requests for data published here should be submitted to citing this study. The authors declare no conflict of interests.

